# The intrinsic clock of the hippocampal subfield CA3 rescues limbic seizures in a biohybrid graft-host interaction in vitro

**DOI:** 10.1101/2023.01.26.525630

**Authors:** Davide Caron, Stefano Buccelli, Ángel Canal-Alonso, Marco Hernández, Giacomo Pruzzo, Juan Manuel Corchado, Michela Chiappalone, Gabriella Panuccio

## Abstract

Hippocampal dysfunction is the hallmark of mesial temporal lobe epilepsy (MTLE), the most common epileptic syndrome in adults and the most often refractory to medical therapy. Deep brain stimulation (DBS) may ameliorate drug-refractory MTLE, but it still cannot guarantee a seizure-free life. One major drawback is that the stimulation policy is informed by trial-and-error rather than by the operating mode of the brain. Thus, optimizing DBS parameters is still an unmet clinical need.

Here, we propose the deployment of hippocampal interictal activity in a biohybrid approach to control limbic ictogenesis. Specifically, an electronic bridge establishes a graft-host interaction between the hippocampal subfield CA3 (graft) and the parahippocampal cortex (CTX – host) of distinct rodent brain slices, both treated with 4-aminopyridine; the electronic bridge relays the graft interictal events to the host via electrical pulses. We show that interictal activity generated by the graft CA3 controls limbic ictogenesis in the host CTX even in the absence of feedback from it, thus likely reflecting an intrinsic anti-ictogenic clock of this brain region.

This work opens a translational perspective for MTLE treatment via biohybrid neuroprostheses relying on the intrinsic clock of incorporated hippocampal cells.

## INTRODUCTION

Epilepsy is a life-threatening progressive neurological disorder characterized by sudden uncontrolled activity of the brain (seizure)^1^. Epilepsy carries among the highest global burden of disease^2^ which is greatly contributed by the poor responsiveness of epileptic patients to currently available treatments: as of today, one out of three epileptic patients keeps experiencing seizures despite anti-epileptic drug polytherapy^3,4^. Among epileptic syndromes, mesial temporal lobe epilepsy (MTLE) is the least responsive to medications and the most frequently diagnosed in the adult population^3^, providing a major contribution to the global burden of epilepsy. While drug-refractory MTLE patients may benefit of electrical deep brain stimulation (DBS), the therapeutic outcome of DBS still suffers for inter-individual variability, ranging from poor efficacy to seizure freedom^5-8^. One major drawback of current DBS is the lack of a unifying framework to establish its parameters, which are still obtained by trial-and-error and based on arbitrary stimulation frequencies. Thus, epilepsy treatment remains an unmet clinical need challenged by the progressive nature of the disorder and its patient-specific clinical features.

In this scenario, biohybrid implantable devices are an emerging concept in the biomedical field for their potential to overcome the drawbacks of current neuromodulation treatments. These devices rely on the combined use of biological components (here, brain cells) and engineered devices, offering the unique feature of exploiting the electrical pattern of biological grafts in a controlled setting^9^. To this end, the ideal biological component would be a cellular population endowed with an intrinsic electrical clock and thus offering a pacemaker-like functionality. In this regard, the hippocampus has been shown to exhibit such features^10,11^; further, in vitro studies have demonstrated the anti-ictogenic properties of the interictal pattern intrinsically generated by the hippocampal subfield CA3 when challenged with convulsant treatments^12-16^. Thus, hippocampal cells may represent a suitable candidate for biohybrid implantable devices for MTLE treatment.

Building upon these concepts, we argued that the hippocampal clock in itself may exert an anti-ictogenic effect, for which hippocampal cells might be deployed in biohybrid neuroprostheses for neuromodulation to prevent seizures. Aiming at the ultimate vision of a bidirectional communication between the host brain and the biohybrid device, the very first step is evaluating the feasibility and efficacy of the biohybrid approach in a unidirectional operating mode. To this end, we have established a unidirectional graft-to-host interaction via an electronic bridge between the hippocampal subfield CA3 (graft) and the parahippocampal cortex (CTX – host) of distinct rodent brain slices, both treated with 4-aminopyridine (4AP). The 4AP-treated CA3 generates interictal events only^10^, whereas the 4AP-treated host CTX generates recurrent ictal discharges^16^; the electronic bridge relays the graft interictal events to the host via electrical pulses, making up a unidirectional biohybrid neuromodulator aimed at preventing ictogenesis in the host (**Figure 1**). We hereafter refer to this biohybrid construct as ‘the biohybrid’. We show that the biohybrid controls limbic ictogenesis in the host CTX even in the absence of feedback from it, thus likely reflecting an intrinsic anti-ictogenic clock of its biological component, i.e., the hippocampal subfield CA3. Our work lays the foundation for biohybrid neuroprostheses relying on the intrinsic clock of incorporated hippocampal cells for MTLE treatment.

**Figure 1.**
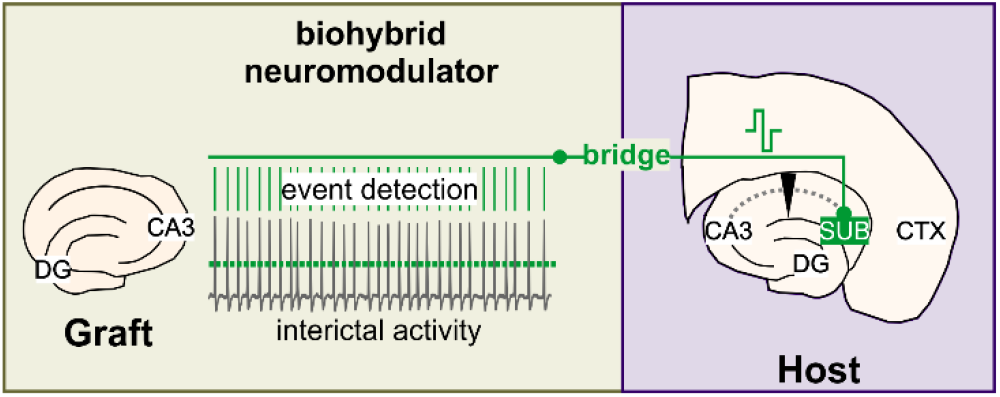
Biohybrid graft-host interaction concept.

## METHODS

### Brain slice preparation and maintenance

Horizontal hippocampus-cortex (CTX) slices, 400 μm thick, were prepared from male CD1 mice 4-8 weeks old. Animals were decapitated under deep isoflurane anesthesia, their brain was removed within 60 s, and immediately placed into ice-cold (∼2 °C) sucrose-based artificial cerebrospinal fluid (sucrose-ACSF) composed of (mM): Sucrose 208, KCl 2, KH_2_PO_4_ 1.25, MgCl_2_ 5, MgSO_4_, CaCl_2_ 0.5, D-glucose 10, NaHCO_3_ 26, L-Ascorbic Acid 1, Pyruvic Acid 3. The brain was let chill for ∼2 min before slicing in ice-cold sucrose-ACSF using a vibratome (Leica VT1000S, Leica, Germany). Brain slices were immediately transferred to a submerged holding chamber containing room-temperature holding ACSF composed of (mM): NaCl 115, KCl 2, KH_2_PO_4_, 1.25, MgSO_4_ 1.3, CaCl_2_ 2, D-glucose 25, NaHCO_3_ 26, L-Ascorbic Acid 1. After at least 60 minutes recovery, individual slices were transferred to a submerged incubating chamber containing warm (∼32 °C) holding ACSF for 20-30 minutes (pre-warming) and subsequently incubated in warm ACSF containing the K^+^ channel blocker 4-aminopyridine (4AP, 250 μM), in which MgSO_4_ concentration was lowered to 1 mM (4AP-ACSF, *cf*. ^17^). All brain slices were incubated in 4AP-ACSF for at least 60 minutes before beginning any recording session. All solutions were constantly equilibrated at pH = ∼7.35 with 95% O_2_ / 5% CO_2_ gas mixture (carbogen) and had an osmolality of 300-305 mOsm/Kg. Chemicals were acquired from Sigma-Aldrich.

For brain slices acting as host, the CA3 was disconnected from the CTX via Schaffer Collaterals disruption. This was either inherent to the slicing procedure or achieved manually by knife-cut. Disconnection was always verified during electrophysiological recording as the lack of propagation of the CA3-driven interictal discharges to the CTX and the persistence of ictal activity in the latter^14,16^.

For brain slices used as graft, the CA3 was isolated by dissecting the CTX out via knife cut; to this end, the blade was placed approximately orthogonal to the Schaffer collaterals. The resulting isolated tissue block comprised the CA3 and dentate gyrus (DG), with only a small portion of the CA1 and the parahippocampal CTX, lateral to the CA3-DG complex. The residual CTX tissue was not connected to the CA3-DG, since both perforant and temporo-ammonic pathways were ablated.

### Microelectrode array recording

Extracellular field potentials were acquired through a 6 × 10 planar MEA (Ti-iR electrodes, diameter 30 μm, inter-electrode distance 500 μm, impedance < 100 kΩ), in which a custom-made low-volume (∼500 μl) recording chamber replaced the default MEA ring (*cf*. ^17^) to attain a laminar flow. Individual slices were quickly transferred onto the recording chamber, where they were held down by a custom-made stainless steel/nylon mesh anchor. Slices were continuously perfused at ∼1 ml/min with 4AP-ACSF, equilibrated with carbogen. Recordings were performed at 32° C, achieved with the use of a heating canula (PH01) inserted at the recording chamber inlet port (temperature set at 37° C) along with mild warming of the MEA amplifier base (temperature set at 32° C), both connected to a TC02 thermostat. The recording bath temperature was checked using a k-type thermocouple. Signal were acquired with the MEA1060 amplifier using the Mc_Rack software or with the MEA2100-mini using the Multichannel Experimenter software, sampled at 2, 5 or 10 kHz and low-passed at half the sampling frequency before digitization. In all experiments, signals were stored in the computer hard drive for offline analysis. The custom recording chamber was designed and made by Crisel Instruments, Italy. The equipment for MEA recording and temperature control was purchased from Multichannel Systems (MCS), Reutlingen, Germany.

### Closed-loop architectures

For n = 5 experiments, the closed-loop architecture was designed in Simulink environment (Mathworks, Natick, USA) receiving the MEA signals through a PCI-6255 DAQ board (National Instruments, USA), and interfaced with the MEA1060 system by a custom PCB (PCB Project, Segromigno Monte, Lucca, Italy). Pulses were pre-programmed in the MC_Stimulus II software and downloaded to the STG2004 stimulus generator (both from MCS), which was triggered via TTL. **Figure 2a** shows the overall closed-loop architecture, whereas **Figure 2b** illustrates the Simulink model. The *Analog input* block reads the MEA signals through the PCI-6255 DAQ board and feeds the CA3 signal from one selected electrode to the *Event detection* block. Detection of the CA3-driven interictal events is based on hard-threshold crossing of the signal amplitude, set manually for each experiment as closely as possible to the signal baseline and no higher than one half of the maximal signal amplitude. The model also implements an *Enabling condition* for triggering the *Stimulation* block, to avoid multiple detections of the same event and to blank the stimulus artifact that would otherwise lead to a false event detection. The enabling condition consisted of a 250 ms window from the last delivered pulse; the window duration was based on prior observation of the minimum interval of occurrence and duration of CA3-driven events and the maximum time-lag and duration of the stimulus artifact. Once triggered, the *Stimulation* block output, in turn, triggers the *Digital Output* block, which sends a TTL signal to the custom PCB controlling the external stimulator (STG2004, MCS). The *Scope* visualizes the MEA signals and the output of the *Event detection, Time since last event*, and *Stimulation* blocks.

**Figure 2.**
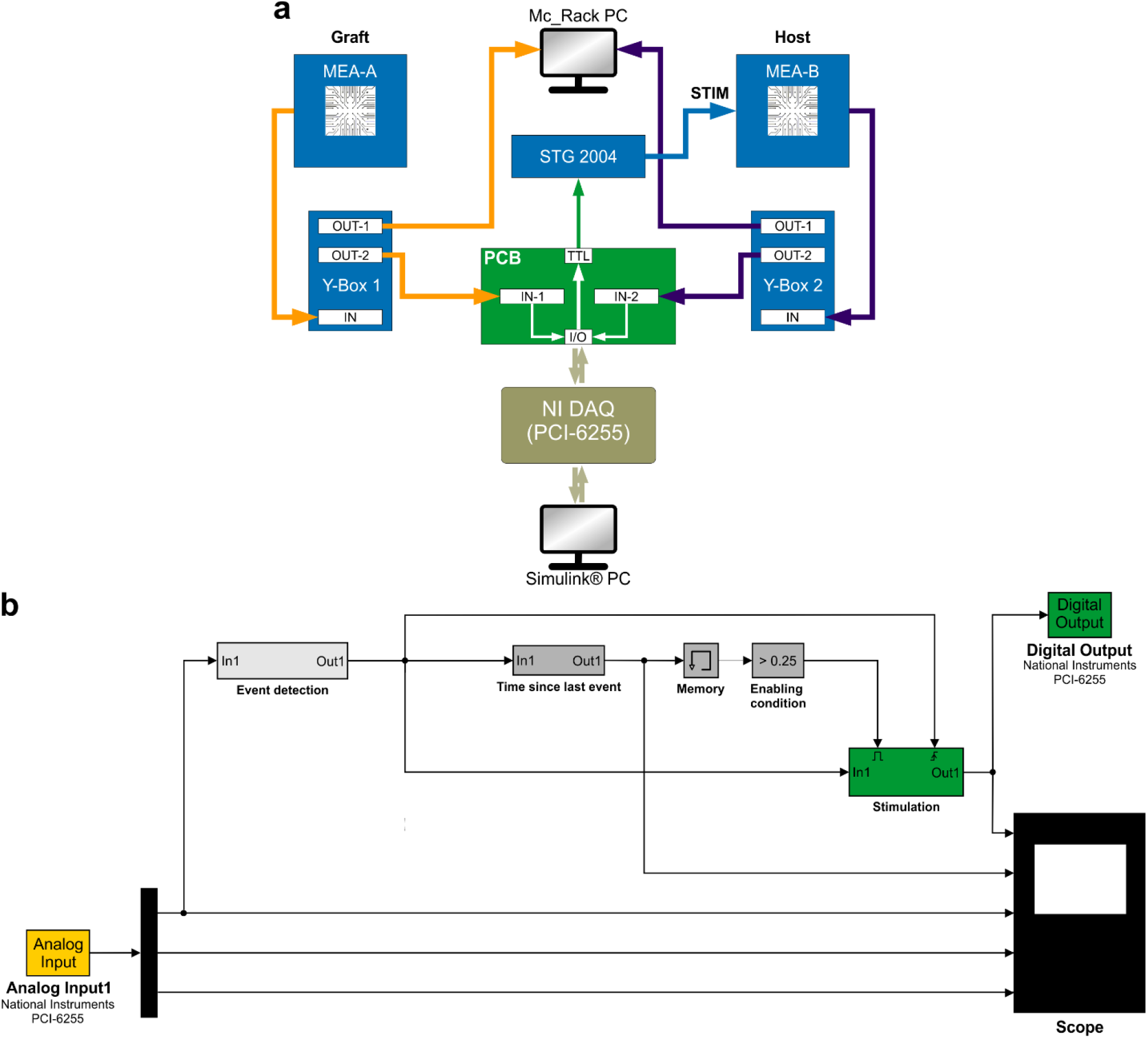
Closed-loop architecture for biohybrid graft-host interaction. **a**. Schematic diagram of the double-MEA closed-loop set-up used to establish the unidirectional graft-host bridge. The signal from each headstage accommodating the graft and the host brain slice is first sent to a signal splitter (Y-box, MCS) which forwards it both to the recording PC (Mc_Rack PC) and to a custom PCB. The latter routes the signals to the National Instruments Digital Acquisition system (NI DAQ-6255) enabling their processing in Simulink environment. The NI DAQ also governs the TTL output of the custom PCB, which triggers the stimulator (STG 2004, MCS) upon detection of an interictal event in the graft CA3. **b**. Simulink model governing the closed-loop architecture, consisting of input (yellow), processing (light grey) and output (green) blocks, along with a scope window (black). The *Analog Input* block retrieves the signals from the MEA system and feeds the graft signal to the *Event detection* block. The latter implements a hard-threshold crossing algorithm and feeds its output to the *Time since last event* block to start the decision routine. The *Stimulation* block is triggered if the enabling condition is met (>250 ms must have passed since the last threshold crossing to avoid multiple detection of the same event as well as false detections due to the stimulus artifact). The enabled *Stimulation* block triggers the *Digital Output* block, which, in turn, activates the TTL signal to the external stimulator. The Scope window enables the real-time visualization of the MEA signals, as well as the output of the *Event detection, Time since last event*, and *Stimulation* blocks.

For n = 3 experiments, the 2-MEA closed-loop architecture relied on the built-in real time feedback (RTF) generator of the MEA2100-mini system. The set-up comprised two Interface Boards (IFB) and two Signal collector Units (SCU), each connected to an MEA60-mini headstage. The stimulation pulse was pre-programmed in the Multichannel Experimenter software and triggered by the graft sub-system via RTF-enabled Digital Output connected to the Digital Input of the host subsystem.

As the closed-loop architecture operated in real-time, the delay between interictal event detection and pulse delivery was equal to the sampling interval.

### Closed-loop graft-host interaction

Stimulation was delivered in the subiculum of the host and consisted of square biphasic direct current pulses (±150-300 μA, balanced charge, 100 μs/phase, positive phase first) conveyed *via* MEA electrode pairs connected in bipolar configuration. A fast input/output (I/O) curve was always performed prior to the first stimulation protocol to find the current amplitude that would reliably elicit an interictal-like discharge in the host (≥80% response rate, *cf*. ^17^). The best performing stimulus amplitude was then kept constant throughout the experiment. Simulation session lasted 20 minutes (> 3 times the mean interval of occurrence of ictal discharges, as observed from the preceding control phase), except for one experiment, where it had to be aborted 1 minute prior (*cf*. note in Table 1). Baseline activity in the absence of stimulation was always recorded before and after each stimulation protocol and is overall referred to as control (CTRL).

**Table 1.** Overview of the experimental dataset and results.

| Mouse n. | Host slice n. | Graft Slice n. | Stimulation parameters |  |  |  |  |  | % ictal state [% reduction] |  |  | Evoked response probability |
| --- | --- | --- | --- | --- | --- | --- | --- | --- | --- | --- | --- | --- |
| | | | $\mu$ | $\sigma$ | CV | duration [s] | n. pulses | pulse amplitude [μA] | CTRL1 | Bridge | CTRL2 | |
| 1 | 1 | 1 | 5.08 | 1.51 | 0.3 | 1200 | 214 | 250 | 19.28 | 0 [100] | 9.32 | 1 |
| 2 | 2 | 2 | 1.02 | 1.16 | 1.14 | 1200 | 1162 | 250 | 15.14 | 3.9 [74.2] | 10.33 | 0.26 |
|  | 3 | 3 | 1.07 | 1.32 | 1.24 | 1140* | 1024 | 300 | 14.95 | 3.6 [75.6] | 17.77 | 0.22 |
| 3 | 4 | 4 | 3.54 | 1.56 | 0.41 | 1200 | 315 | 150 | 7.66 | 0 [100] | 9.41 | 0.73 |
|  | 5 |  | 6.52 | 1.49 | 0.23 | 1200 | 171 | 200 | 8.22 | 0 [100] | 6.8 | 0.98 |
| 4 | 6 | 5 | 0.97 | 1.14 | 1.18 | 1200 | 1233 | 225 | 18.82 | 0 [100] | 12.74 | 0.49 |
| 5 | 7 | 6 | 2.07 | 1.99 | 0.96 | 1200 | 487 | 150 | 24.58 | 0 [100] | 31.57 | 0.94 |
| 6 | 8 | 7 | 1.51 | 1.18 | 0.78 | 1200 | 783 | 350 | 24.76 | 0 [100] | 17.3 | 0.48 |

\*Bridge aborted 60 s before the end of the established duration because of technical issues with the graft perfusion.

### Data analysis

Ictal events were labeled semi-automatically using custom software written in MATLAB R2021b (MathWorks, Natick, USA), described in^18^, and manually corrected as required. Quantification of ictal activity was then used to compute the time % spent in the ictal state (*P*_*ictal*_) with respect to the observation time window (*t*_*TOT*_), which was kept similar to the stimulation duration for reliability. The *P*_*ictal*_ was computed as per Equation 1:

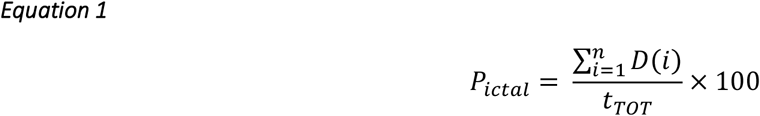

where *i* is i^th^ ictal discharge, *D* is the ictal discharge duration and *t*_*TOT*_ is the total observation time. For CTRL phases (no stimulation), *t*_*TOT*_ is the time between the onset of the first and the termination of the last measured ictal event; for stimulation phases (STIM), *t*_*TOT*_ is the duration of the stimulation protocol.

Interictal-driven cortical responses were also labeled semi-automatically. To this end, the pulse timings stored in the recording were used to remove the stimulus artifact; this was attained by replacing a post-stimulus window of 5 ms with zeros. The evoked responses where then labeled with a peak clustering algorithm and manually corrected as required. Then, the evoked response probability (*P*_*evoked*_) was computed as in **Equation 2**:

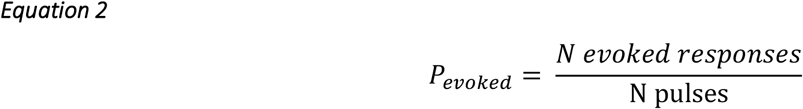

Fitting of the μ vs CV relationship was performed in MATLAB R2021b using the mono-exponential function in **Equation 3**:

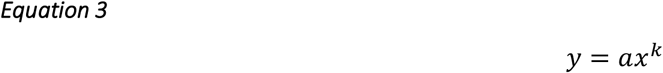

Throughout the text, data are reported as mean ± SEM. For the *P*_*ictal*_, the mean is expressed as arithmetic mean ± SEM. For pulse distribution parameters, we computed the geometric mean (μ) and standard deviation (σ), since the inter-pulse intervals were lognormally distributed (*cf*. ^*18*^); the coefficient of variation (CV) was computed as σ/μ.

### System impulse response estimation

To compare the impulse response of cortical networks during biohybrid stimulation with those driven by the CA3 interictal events in the intact hippocampal loop, we computed the exponential time constant (τ) of cortical events in the two scenarios. To this end, we considered the field potentials generated by brain slices as a linear time-invariant system characterized by **Equation 4**:

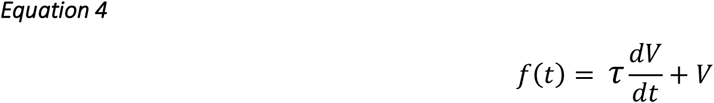

A solution to **Equation 4** for initial value *V*_*0*_ is shown in **Equation 5**:

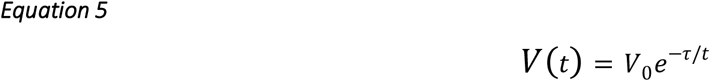

The system decays to baseline when *V*(*t*) ≈ 0 (the value cannot be considered exactly 0 because of the natural oscillations of a signal). To calculate the value of τ, we first zeroed the signal baseline by subtracting the baseline mean from the signal, then used the signal natural logarithm to obtain ***lnV(t)*** and ***t***, and considered *V*_0_ as the peak of the impulse.

### Statistical analysis

Statistical analysis was performed using SPSS (IBM, Armonk, NY, USA). The efficacy of the biohybrid was statistically assessed using ANOVA followed by the Games-Howell post-hoc test. The latter was chosen based on normality (Shapiro-Wilk test) and homoskedasticity (Levene’s test) check. For comparison of the impulse response, we first transformed the obtained τ values into a normal distribution using the Box Cox transformation; then, we computed the average values for each experiment and compared the two datasets using the Welch’s test. Throughout the text, data are expressed as mean ± SEM, unless otherwise specified. Results statistics were considered significant for p < 0.05.

## RESULTS

### The biohybrid controls limbic ictogenesis

The experimental dataset comprises n = 8 host brain slices and n = 7 graft brain slices from 6 mice. Graft brain slices were randomly selected without any a priori consideration about the statistics of their interictal pattern. This yielded a population of grafts exhibiting variable mean (μ) and standard deviation (σ) of their interictal event intervals. **Figure 3a** shows a representative experiment in which the biohybrid completely prevented ictal activity in the host brain slice. For simplicity, the CTRLs show the signal recorded from the host CTX only. The representative trace segment during biohybrid stimulation illustrates its time-locked operation with the biological component (graft CA3, purple) and the responses driven in the host CTX (black); the pulse timings are marked by the green bars. As summarized in **Figure 3b**, the biohybrid significantly decreased the percentage of ictal state (CTRL1: 16.68 ± 2.46; biohybrid: 0.94 ± 0.66; CTRL2: 14.40 ± 3.01; one-way ANOVA, F(df): 15.98(2); bridge vs CTRL1: p < 0.0005; biohybrid vs CTRL2: p < 0.005; CTRL1 vs CTRL2: p = 0.81). More in detail, the biohybrid fully prevented ictal activity in 6 out of 8 host CTX, whereas in the remaining 2 it decreased it by 74.2% and 75.6%, yielding an overall ictal state reduction of 93.73 ± 4.39%. In all experiments, ictal activity reappeared upon stimulus withdrawal confirming that its disappearance during the stimulation was due to the action of the biohybrid.

**Figure 3.**
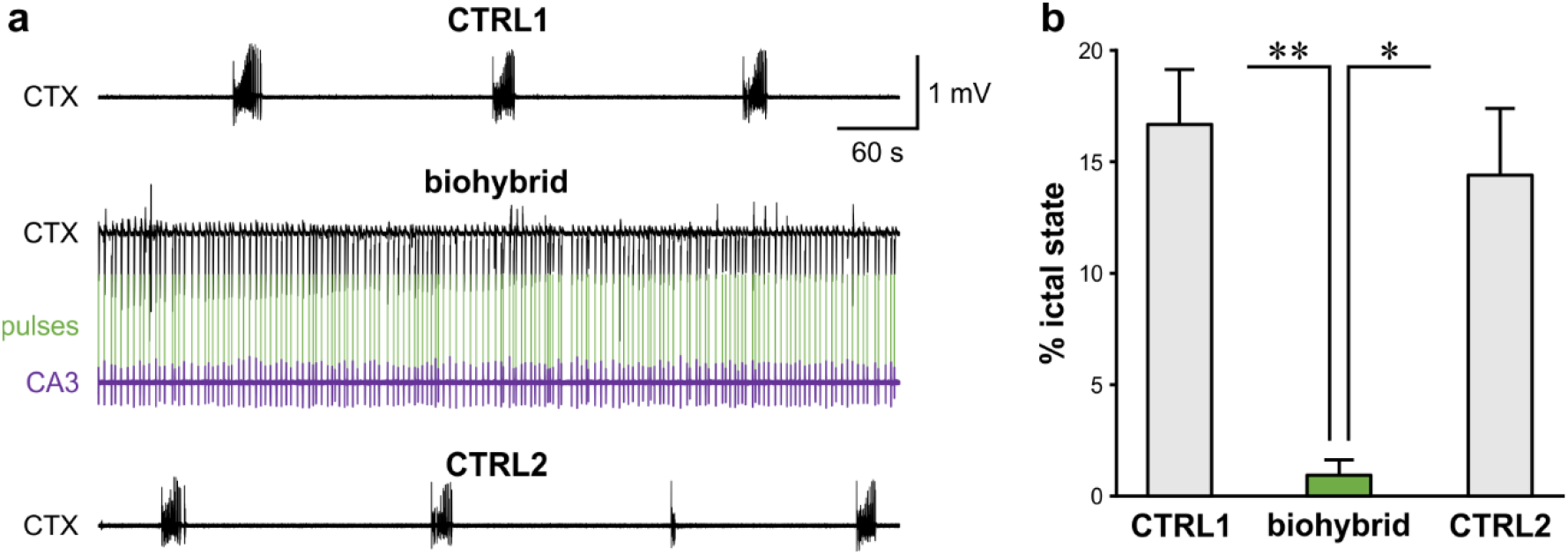
Biohybrid modulation controls parahippocampal CTX ictogenesis. **a**. Representative MEA recording of a best-case experiment showing recurrent ictal discharges in baseline condition (CTRL1), their abolishment by the bridge and their reappearance upon bridge withdrawal (CTRL2). Stimulus artifacts in the bridge signal are partially removed for clarity and are marked by the vertical green bars; the CA3 signal (purple) is shown to demonstrate that electrical pulses are time-locked with the interictal events generated by this brain region. **b**. Summary of the results statistics obtained for n = 8 experiments, showing the significant reduction in ictal state % during the bridge. * p < 0.005; ** p < 0.0005.

### The efficacy of the biohybrid relies on the intrinsic temporal statistics of its biological component

If the efficacy of the biohybrid stems in an intrinsic anti-ictogenic clock of its biological component, the diverse stimuli distributions, reflecting biological variability, should not yield different degrees of ictal activity reduction. **Figure 4a** illustrates the wide range of the mean interval (μ) of the CA3-driven electrical pulses and their coefficient of variation (CV); in keeping with previous findings, the two parameters exhibit a power-law relationship^18^, denoting the existence of a finely regulated intrinsic clock within the CA3 (*cf*. also ^10,11^). As shown in **Figure 4b**, the degree of ictal state reduction did not correlate with the pulse distribution parameters; in n = 6 brain slices (i.e., 75% of the cases) the biohybrid fully prevented ictal activity, regardless of the temporal statistics of the delivered pulses. In the remaining n = 2 brain slices (25% of the cases), the biohybrid still achieved a very good percentage of ictal reduction (∼75%); however, because this paradigm fully prevented ictal activity in the majority of the cases, this result was surprising and it appeared to be in contrast with the majority of the dataset. One possible reason underlying this observation is the poor entrainment of cortical networks by the stimulation. To address this possibility, we evaluated the evoked response probability. As shown in **Figure 4c**, this analysis unveiled the poor responsiveness of the brain slices in which the biohybrid did not fully prevent ictal activity (probability = 0.22 and 0.26); in contrast, brain slices in which ictal activity was fully prevented exhibited a higher responsiveness to stimulation, with an evoked response probability ranging from 0.48 to 1.

These findings suggest the presence of an intrinsic anti-ictogenic clock within the CA3. Further, the probability of entraining cortical networks by biohybrid stimulation might reflect the timing of stimulation relative to the cortical state within the interictal-to-ictal transition.

To further address these possibilities, we estimated the impulse response of the CTX during biohybrid stimulation and compared it to that of cortical networks responding to the CA3 biological inputs within the intact hippocampal loop (n = 8 brain slices for each experimental group). The impulse response is the output reaction of a dynamical system to an external perturbation and it decays as a function of time dependent on the system time constant (τ). The latter can thus be used to address if the system is back to a relaxation state before the next impulse. In the case of electrical stimulation, the value of τ relative to the pulse spacing may influence the probability of evoking a response and thus of entraining the stimulated brain network.

**Figure 5** shows representative signals and the measured τ values from the CTX within the intact hippocampal loop **(a)** and during biohybrid stimulation **(b)**. As expected, the τ values changed in time, reflecting the dynamical behavior of the CTX along the succession of the received inputs. Further, each brain slice exhibited a different relationship between the input intervals and the τ values, both in the intact loop (**Figure 5c**) and during biohybrid stimulation (**Figure 5d**). However, as shown in **Figure 5e**, the obtained τ values were significantly faster during biohybrid stimulation (median τ values range – intact hippocampal loop: 0.07 - 0.2 s; biohybrid: 0.002 - 0.04 s; n = 8 brain slices each group; p < 0.001; Welch test).

These results are consistent with the inter-individual variability of biological systems, and suggest that the biohybrid induces an adaptive behavior specific to each host CTX; the impulse response of the latter may thus be state-dependent. The faster impulse response during stimulation may reflect the inherent difference between a biological input and its artificial surrogate (electrical pulse), as well as the lack of CTX-to-CA3 network reverberations seen in the intact hippocampal loop.

**Table 1** provides an overview of the experimental dataset, the used stimulation parameters, and the performance of the biohybrid. Note that graft n. 4 was deployed in two host brain slices (n. 4 and 5), and that the statistical parameters of its interictal events were different in the two cases; yet, the biohybrid achieved 100% prevention of ictal activity in both hosts, further substantiating the intrinsic anti-ictogenic clock of the CA3 and the inherent adaptive behavior induced by the biohybrid in the host. In turn, these features enable the unsupervised operation of the biohybrid, as opposed to current DBS policies defined by the human operator.

Taken together, these results suggest that the intrinsic clock of hippocampal cells may be deployed in a biohybrid implantable device for MTLE treatment; closing the loop in the other direction to attain a bidirectional communication bridge is expected to optimize the device operation to achieve a higher degree of host network entrainment and thus of seizure prevention.

## DISCUSSION

We have presented the proof-of-concept of a biohybrid neuromodulator for MTLE treatment based on the intrinsic anti-ictogenic clock of the hippocampal subfield CA3. Previous work has demonstrated that the CA3 is the site of origin of a fast interictal pattern^10,14^ which acts as pacemaker-like driver of synchronicity^10,11^ and serves an anti-ictogenic function^12-16^. Building upon this, more recent work has corroborated the concept of intrinsic anti-ictogenic effect of the CA3-driven interictal pattern by deploying open-loop stimulation surrogating its temporal dynamics to control limbic ictal activity^18^; this work has also demonstrated the power-law relationship linking the mean (μ) and coefficient of variation (CV) of the interictal event intervals, suggesting the existence of a finely tuned intrinsic clock within the CA3. Here, we have pushed forward the concept of the CA3 as an anti-ictogenic neuromodulator by devising a biohybrid paradigm for limbic seizure control. In our hands, the biohybrid exhibited an outstanding performance, with full prevention of ictal activity in 75% of the host CTX slices. Remarkably, the stimulation pattern did not correlate with the degree of efficacy of the biohybrid (*cf*. **Figure 4**), corroborating the view of a finely tuned anti-ictogenic clock within the CA3.

**Figure 4.**
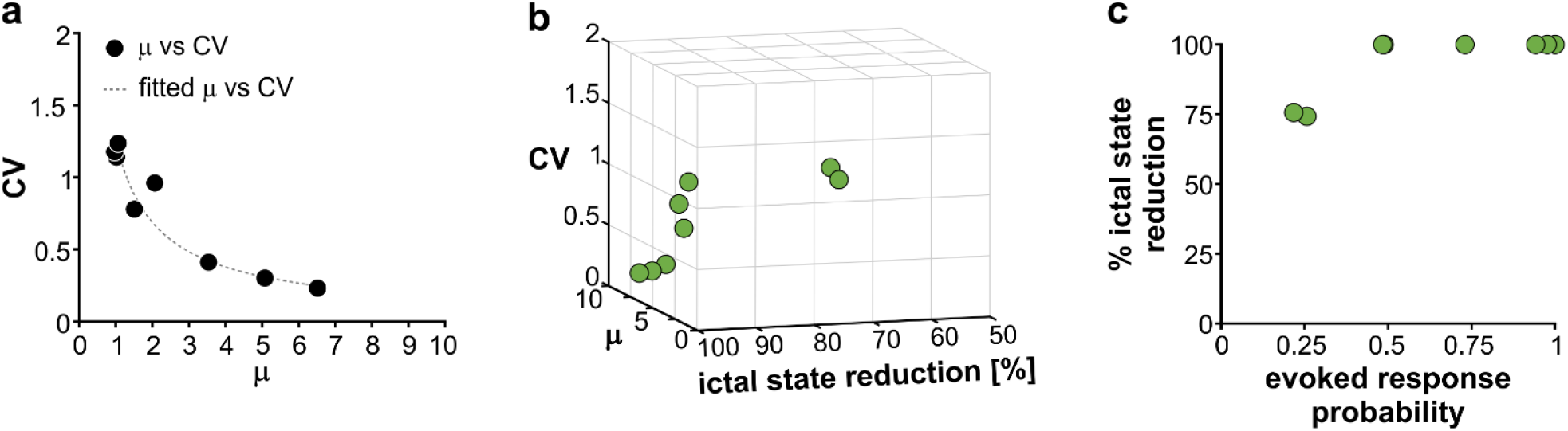
The efficacy of the biohybrid relies on the intrinsic clock of the graft CA3. **a**. The statistical parameters μ and CV describing the temporal distribution of the CA3-driven pulses exhibit a variable range and their relationship follows a power law. **b**. 3D plot showing the relationship among μ, CV and % of ictal state reduction. The efficacy of the biohybrid is not bound to specific stimuli distribution parameters. **c**. Probability of cortical evoked responses vs the degree of ictal state reduction. The efficacy of the biohybrid reflects the effective entrainment of cortical networks by electrical stimulation.

**Figure 5.**
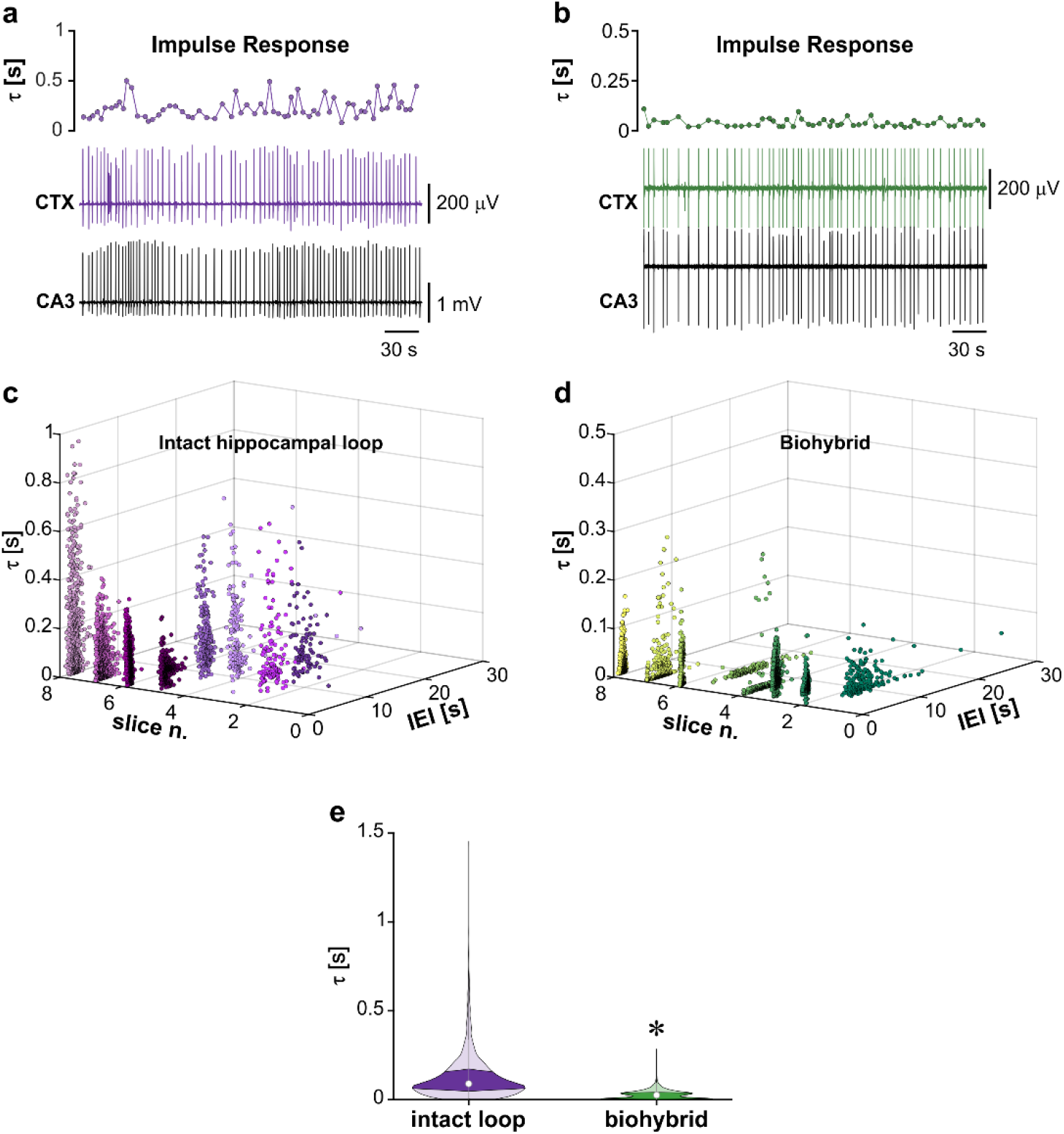
Impulse response of cortical networks within the intact hippocampal loop and during biohybrid stimulation. **a**. Representative MEA recording from the CA3 and CTX within the intact hippocampal loop, and corresponding τ values, showing the time-varying impulse response of cortical networks to the CA3 biological inputs. **b**. Representative MEA recording from the CA3 and CTX during biohybrid stimulation; similar to the intact hippocampal loop, the cortical impulse response exhibits a pulse-to-pulse variability. Note that the τ values are smaller than what measured in the intact hippocampal loop. **c-d**. Relationship between interictal event interval (IEI) and impulse response for each brain slice within the intact hippocampal loop **(c)** and biohybrid stimulation **(d)** datasets. Inter-individual variability is consistently observed in both scenarios. **e**. Biohybrid stimulation indices faster impulse responses than what measured in the intact hippocampal loop. * p < 0.005.

A more in-depth analysis unveiled that the 25% of host CTX slices in which the biohybrid did not fully prevent ictal activity exhibited a poor entrainment by electrical stimuli, with a very low probability of the evoked responses (∼0.25). However, a fast I/O was always performed prior to the stimulation session to address the preservation of subicular projections to the CTX and to find the appropriate stimulus intensity; thus, we may hypothesize that the timing of stimulation relative to the time course of the interictal-to-ictal transition plays a role in the responsiveness of brain networks to electrical stimulation. In support of this view, we have shown that cortical networks exhibit dynamical impulse responses both in the intact hippocampal loop and during biohybrid stimulation. This view is also supported by recent evidence ascribing the pro-or anti-ictogenic role of interictal activity (and external perturbations in a broader perspective) to epileptic networks bistability, making them more prone to generate seizures in response to external perturbations when they are in a specific state transition^19^. In keeping with this, it has been demonstrated that the seizure-propensity of brain networks is preceded by their loss of resilience to perturbations^20^. In this regard, a bidirectional communication bridge closing the loop in the other direction (i.e., from the host back to the biohybrid) will likely achieve a higher degree of host brain network entrainment (higher evoked response probability), as expected from a process of dynamic mutual adaptation; in turn, this is expected to achieve a higher degree of seizure prevention. By demonstrating the unidirectional operating mode of the biohybrid we have made the very first step toward its deployment in a bidirectional bridge. Thus, our work brings a substantial conceptual difference with open-loop stimulation informed by the temporal statistics of the CA3-driven interictal pattern^18^.

Biohybrid designs are an emerging concept in the biomedical field for brain implantable devices^9^. Their primary advantage stems in deploying biological information processing via cellular elements to sense/control brain function, thus bypassing the trial-and-error search of stimulation parameters. As opposed to regenerative approaches, the cellular component making up the biohybrid device is not directly grafted into the host brain, hence its behavior can be finely and safely controlled. Thus, biohybrid brain implants offer the unique advantage of biological vs hardware or software signal processing, but without the potential risks of cell grafting.

The faster impulse response of the CTX during biohybrid stimulation may reflect the inherent difference between a biological input and its artificial surrogate (electrical pulse): while biological inputs vary in amplitude, duration, and phase of the target brain region when the input reaches it, electrical pulses are delivered at a constant amplitude and duration, and the sole varying parameter is the phase of the stimulated brain network at the time of stimulation. Further, the longer impulse response of cortical networks within the intact hippocampal loop might reflect the CTX-to-CA3 network reverberations, which were inherently absent in the experimental setting of the present work. In turn, this further supports the importance of implementing the directed connectivity of the host brain neurons to the biological component of the biohybrid device. Microfluidics-based biological micro-electro-mechanical systems (bio-MEMS) would enable such biohybrid approach^21^.

As future directions, there are several undertakings that would contribute significantly to push forward the concept of biohybrid neuromodulation: **(i)** closing the loop to attain a bidirectional biohybrid interaction, **(ii)** deploying isolated hippocampal neurons (as opposed to isolated CA3 sections) to step closer to the biohybrid device implementation, **(iii)** challenging the biohybrid against other epilepsies than MTLE, and, ultimately **(iv)** devising the bio-MEMS for in vivo deployment toward its clinical translation.

## DECLARATION OF INTERESTS

The authors have no competing interests to declare.

## ETHICS STATEMENT

All procedures were approved by the Institutional Animal Welfare Body and by the Italian Ministry of Health (authorizations 860/2015-PR and 176AA.NTN9), in accordance with the National Legislation (D.Lgs. 26/2014) and the European Directive 2010/63/EU. All efforts were made to minimize the number of animals used and their suffering.

## ACKNOWLEDGEMENTS

This work was funded by the European Union under the Horizon 2020 framework program: H2020-MSCA-IF-2014 Re.B.Us – Rewiring Brain Units, GA n. 660689 awarded to GP; H2020-FETPROACT-01-2018 (RIA) HERMES – Hybrid Enhanced Regenerative Medicine Systems, GA n. 824164. We thank Ms. Alessandra Sanna and Ms. Cinzia Nasso for administrative assistance. We also thank the support of PCB project (Segromigno Monte, Lucca, Italy) for the custom PCB used in the experiments reported in this manuscript.

## CRediT AUTHORSHIP CONTRIBUTION STATEMENT

**Davide Caron:** formal analysis, investigation, validation, writing – review & editing. **Stefano Buccelli:** investigation, methodology, software, validation, writing – review & editing. **Ángel Canal-Alonso:** formal analysis, methodology, software, validation, writing – review & editing. **Marco Hernández:** formal analysis, validation, writing – review & editing. **Giacomo Pruzzo:** conceptualization – hardware, writing – review & editing. **Juan Manuel Corchado:** formal analysis, validation, resources, supervision, writing – review & editing, funding acquisition. **Michela Chiappalone:** conceptualization, funding acquisition, methodology, project administration, resources, supervision, validation, writing – review & editing. **Gabriella Panuccio:** conceptualization, data curation, formal analysis, funding acquisition, investigation, methodology, project administration, resources, software, supervision, validation, visualization, writing – original draft, review & editing.

## Notes

### Competing Interest Statement

The authors have declared no competing interest.

